# Personality assessment of synanthropic rhesus macaques: implications and challenges

**DOI:** 10.1101/2024.07.30.605931

**Authors:** Taniya Gill, Anshul Gautam, Jorg J.M. Massen, Debottam Bhattacharjee

## Abstract

“What makes animals thrive in human-dominated environments?” is a question that has been extensively researched transcending disciplines, but findings remain inconclusive. Consistent inter-individual differences or personalities can potentially explain the functional significance of habitat-specific traits and their variations that help animals successfully coexist with humans. Rhesus macaques (*Macaca mulatta*) are the most successful non-human primate in the Anthropocene, living in diverse climatic and environmental conditions. Studying the personalities of synanthropic rhesus macaques can provide insights into the biological traits that facilitate their success in human-dominated environments. We employed a multi-method ‘bottom-up’ approach of behavioral observations and novelty experiments, standardized for assessing captive non-human primates, to evaluate the personalities of synanthropic adult rhesus macaques (*N=52*). To our surprise, novelty experiments encountered significant challenges, limiting their effectiveness. However, behavioral observations in the form of focal sampling revealed two repeatable traits: *sociability* and *cautiousness*. We found an effect of sex on sociability, where males were more sociable than females. In an additional analysis, we found that individuals who obtained food through contact provisioning had higher cautiousness scores than individuals who obtained food through non-contact provisioning. We discuss how the observed personality traits and their variations potentially offer adaptive advantages in human-dominated environments, where rhesus macaques face both benefits, such as anthropogenic subsidies and reduced predation, and costs, like exposure to anthropogenic stressors. We also emphasize that protocols designed for captive conditions may not be directly applicable to free-living animals. Thus, the study underscores the need to reconsider experimental designs to obtain comparable empirical evidence between captive and non-captive populations to enhance the ecological validity of personality assessments. Nevertheless, empirically identifying traits using observations in synanthropic species like rhesus macaques can still provide valuable insights into the mechanisms that enable certain animals to thrive amidst a rapid expansion of anthropogenic activities.

**Research highlights:** Studying personalities of synanthropic animals can shed light on the biological traits that help them thrive in human-dominated environments.

A ‘bottom-up’ multi-method approach, standardized for assessing captive NHP personalities, was employed to study a population of synanthropic rhesus macaques.

Behavioral observations captured two repeatable traits, namely *sociability* and *cautiousness*, where comparatively more cautious individuals obtained food through contact provisioning. Novelty experiments faced significant challenges.

## 1. Introduction

Anthropogenic activities have intensified manifold in recent years, impacting the lives of non-human animals (hereafter, animals) in every possible habitat (Dirzo et al., 2014). Such an increasing expansion has resulted in highly porous boundaries between us and the wildlife around us. Subsequently, this human-wildlife coexistence has garnered exorbitant attention from diverse disciplines, like ecology, behavior, cognition, and even social sciences. From the behavioral ecology perspective, one of the critical questions remains: what makes some animals thrive in human-dominated environments? Although considerable scientific advancements have been made in addressing the question, like the presence of ‘unlimited’ behavioral plasticity in animals (Sih et al., 2004), the answers remain far from conclusive. In contrast to the view of behavioral plasticity, personality or consistent inter-individual differences suggest that behavioral responses are rather ‘non-flexible’ and are somewhat constrained (Réale et at., 2007; Réale et al., 2010). While to what extent biological traits have plasticity or constraints is an ongoing topic of research (Dingemanse et al., 2010), inter-individual differences in the phenotypic expression of specific traits can potentially explain their ‘selective’ advantages with regard to certain environmental conditions and habitats (Santicchia et al., 2018; Belgrad et al., 2018; Leclerc et al., 2016). Due to the strong associations of personality traits with adaptive and survival benefits (Smith & Blumstein, 2008; Moiron et al., 2020), studying these traits in animals living close to humans can provide valuable insights into their potential functional advantages.

The study of animal personality has advanced considerably in the last two decades, encompassing a wide range of taxa, from insects to mammals, including non-human primates (NHPs). In particular, for NHPs, testing environments vary, including captive and free-ranging living conditions, existing social settings, or isolation from social groups. While captive animals can be tested either in existing social settings or in isolation and researchers have discussed the trade-offs (Šlipogor et al., 2016; Koski, 2011; Koski & Burkart, 2015), free-living conditions typically do not allow testing animals in isolation. Besides, for gregarious animals like most NHPs, assessing personality traits in existing social settings may provide benefits of higher ecological relevance (Koski, 2011). In addition to testing environments, there are methodological differences, ranging from subjective questionnaire-based ratings to less subjective (or more objective) observational or experimental approaches (Freeman et al., 2011). The questionnaire-based approach involves rating animals (often) on predetermined traits, primarily by caregivers based on their day-to-day interactions with the animals (Freeman et al., 2013); in contrast, the observational or experimental methods include testing traits that are salient or ‘rare’ using ethograms (Koski, 2011; Freeman et al., 2013; Massen et al., 2013; Šlipogor et al., 2016). While these methods have their advantages and disadvantages (see Vazire et al., 2007; Freeman et al., 2013; Šlipogor et al., 2020), the combined multi-method approach of behavioral observations and experiments is relatively less utilized. Further, it has been highlighted that observations and experiments as standalone methodologies may provide evidence of unrelated and independent personality traits (Martinig et al., 2022). Therefore, a multi-method approach can reveal consistent salient as well as ‘rare’ traits more comprehensively (Kluiver et al., 2022; Bhattacharjee et al., 2024a).

Several NHP species live in and around human-dominated environments, but rhesus macaques are the most successful species in the Anthropocene (Cooper et al., 2022). Rhesus macaques are highly adapted to a wide range of climatic (from temperate to tropical) and environmental (from mountainous and forested to semi-deserts and swamps) conditions (Cooper et al., 2022). Likewise, their geographical distribution spreads across Central, South, and Southeast Asia (Brandon-Jones et al., 2004; Fooden, 2000). The success story of rhesus also includes them being a synanthropic species that benefits from humans by living close to them (Klegarth et al., 2017). Synanthropic animals may benefit through indirect (utilizing anthropogenic subsidies, low to no natural predation pressure, etc.) as well as direct (getting provisioned by humans) ways (McLennan et al., 2017); however, there are costs, too. Synanthropes are exposed to anthropogenic stressors (like noise, conflict with humans, etc.) that can considerably impact their survival (Pascal et al., 2020; Ilhman, 2024). Empirical studies have shown that humans can directly influence the social interaction networks and behavioral dynamics of animals living close to them (Lowry et al., 2013; Kaburu et al., 2019; Bhattacharjee & Bhadra, 2020; Bhattacharjee & Bhadra, 2021; Balasubramaniam et al., 2021). Despite the trade-offs, rhesus macaques, a relatively new species in the evolutionary timescale, have successfully coexisted with humans (Cooper et al., 2022). Therefore, assessing the personalities of these synanthropic rhesus macaques can shed light on the biological traits that potentially help them thrive in human-dominated environments.

Rhesus macaques have been studied for their personalities predominantly using questionnaire-based ratings and observational approaches in socially housed captive and natural free-ranging populations (Weiss et al., 2011; Morton et al., 2013; Sussman et al., 2013; Brent et al., 2014; Adams et al., 2015; von Borell et al., 2016; Kohn et al., 2016; Robinson et al., 2018; Altschul et al., 2019). In questionnaire-based studies, traits like *friendliness* or *sociability*, *confidence*, *dominance*, *openness*, *anxiety*, *equability*, *excitability*, etc., were predominantly identified. On the other hand, observational studies labeled traits as *sociability*, *aggressiveness*, *cautiousness*, *fearfulness*, *meek*, *loner*, *nervous*, *passive*, etc. In addition, personalities were associated with dominance status (Kohn et al., 2016), social style (Sussman et al., 2013; Adam et al., 2015), health and welfare (Robinson et al., 2018), and facial dimensions (Altschul et al., 2019). Unlike the questionnaire-based and observational approaches, studies using experimental and further multi-method approaches (i.e., observations and experiments) are almost non-existent in rhesus macaques. In the current study, we implemented a multi-method ‘bottom-up’ approach standardized for assessing captive NHPs (Kluiver et al., 2022; Bhattacharjee et al., 2024a) to determine the personalities of synanthropic rhesus macaques. The multi-method approach included continuous focal observations and novelty experiments (novel objects, novel food items, food puzzles, predator model exposure, cf. Massen et al., 2013; Kluiver et al., 2022; Bhattacharjee et al., 2024a) conducted in macaques’ existing social and free-living environmental conditions.

Several (comparative) experimental studies implementing similar or identical laboratory protocols to relatively ‘noisy’ field conditions have succeeded despite some uncontrollable challenges (Neumann et al., 2013; Molesti & Majolo, 2016; Morand-Ferron et al., 2016; Brubaker et al., 2017; Shaw & Schmelz, 2017; Brubaker et al., 2019). Therefore, we hypothesize that the standardized multi-method approach of personality assessment (for captive NHPs) would effectively capture both salient behaviors and behaviors of rare occurrences in synanthropic rhesus macaques. Due to the ‘bottom-up’ nature of the methodology, informed predictions on personality traits are challenging to make. However, given the despotic social style and steep linear hierarchies of rhesus macaques (Thierry, 2000), we expect to find traits like *fearfulness*, *cautiousness*, etc., that previous observational approaches also reported. In addition, the matrilineal hierarchical system of rhesus macaques fosters strong social associations among philopatric females (Sterck et al., 1997). Thus, we expect traits like *openness* or *sociability* to be present, with females showing higher openness or sociable tendencies than males. Although synanthropic animals are well-adapted to living in human-dominated environments, a ‘landscape of fear’ can not be disregarded (Gallagher et al., 2017), where traits like *vigilance* or *cautiousness* would be expected. In line with previous findings on other relatively despotic NHPs (Bhattacharjee et al., 2024a), we also predict that traits like *exploration* or *persistence* would be present.

## 2. Materials and Methods

### 2.1. Study site and subjects

We conducted the study at the Kamla Nehru Ridge Biodiversity Park (KNR) (28°40’50“N, 077°13’01“E) in North Delhi, India. KNR is a part of the highly weathered Aravalli hill range and is spread out across an 87-hectare area surrounded by motorable roads and dense urban residential areas. The flat hilltops and relatively shallow valleys give the area an undulating topography. The natural vegetation resembles that of tropical dry and mixed deciduous forests. However, a major biodiversity re-introduction and restoration drive is in place to eradicate ‘invasive’ species like the Mexican weed *Prosopis juliflora*. In KNR, a total of five water bodies are present. People use KNR as a walking park, and the area is also known to attract tourists due to its striking biodiversity profile (Sinha, 2014). Among the fauna, a group of free-living rhesus macaques (estimated population of > 100 individuals) is present (**Figure 1**). However, comprehensive information on the population size and other demographic properties, like age and sex, is unavailable. The macaques are provisioned daily by local people (**Figure S1**), typically between 9 AM and 11 AM (T.G., personal observations, 2021-2022). We randomly selected 52 adult macaques (female = 43, male = 9) for the current study, which lasted between early November 2021 and late July 2022. The first four months (November 2021 – February 2022) included identifying the individuals using facial features and scar marks, as well as habituation of the group to the experimenters.

**Figure 1.**
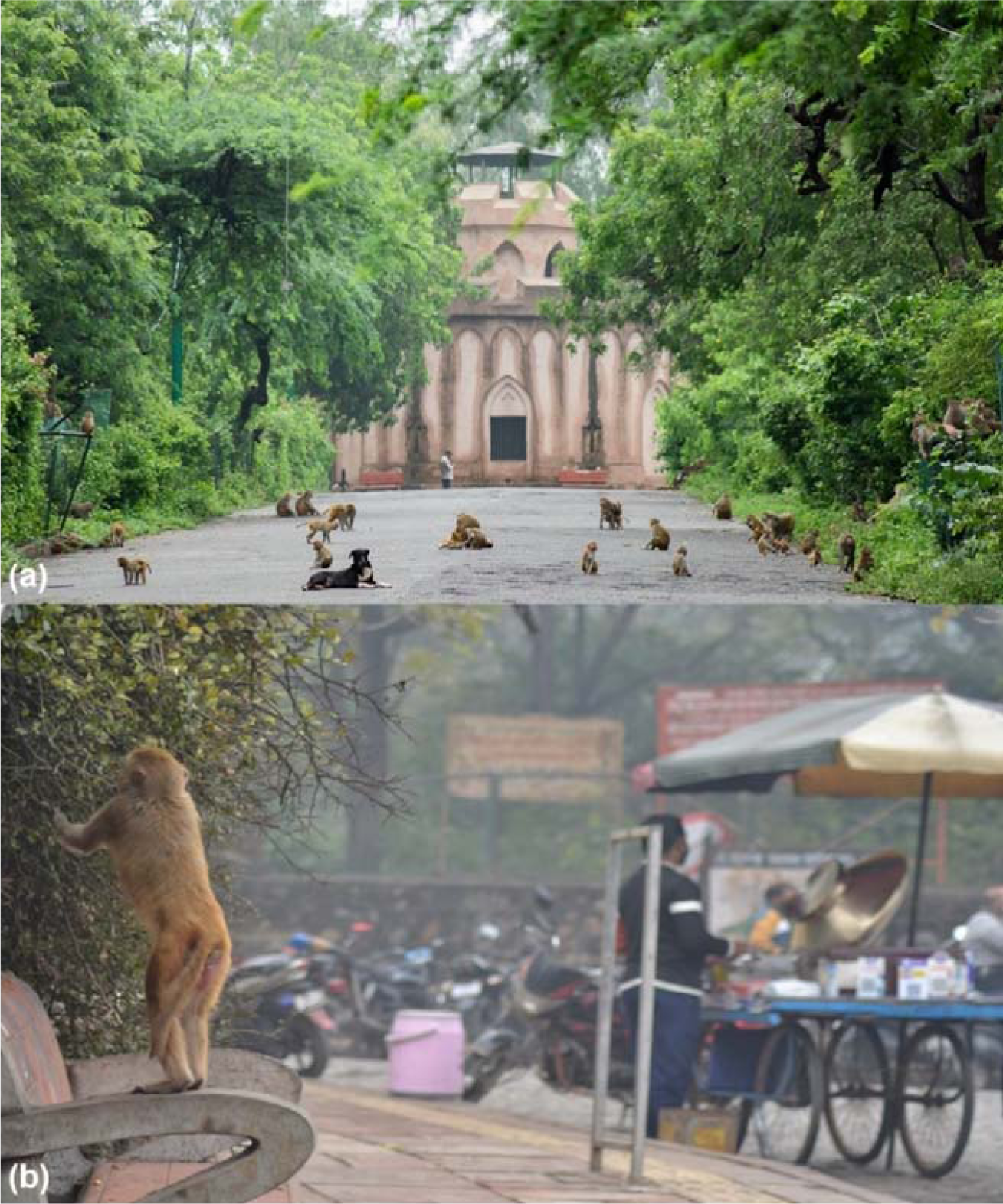
Synanthropic rhesus macaques at the Kamla Nehru Ridge Biodiversity Park, New Delhi, India. *(a)* Some individuals from the study population, along with a free-ranging dog *(b)* A depiction of rhesus macaque presence in human-dominated environments in the study area. [Photo credit: Taniya Gill, 2022]

### 2.2. Data collection for personality assessment

We aimed to use a standardized multi-method bottom-up approach of behavioral observations and novelty experiments to assess personality (Koski, 2011; Massen et al., 2013; Kluiver et al., 2022; Bhattacharjee et al., 2024a). Such a comprehensive approach allows for objectively capturing regular and ‘rare’ behaviors.

#### Behavioral observations

Twenty-minute-long continuous focal observations were conducted following an extensive ethogram (**Table S1**). The ethogram consisted of 87 behaviors, state (durational), and point (frequency) behaviors (which became 110 after including the passive behaviors, i.e., behaviors directed to focal individuals). During focal observations, we also collected data on human-macaque interactions with regard to obtaining food through contact and non-contact provisioning (cf. **Table 1**). Observations were performed five days a week at different times of the day between 9 AM and 5 PM using a pseudo-randomized order, such that an individual was not observed more than once on the same day or on two consecutive days. After correcting for the time individuals were out of sight, a total of 5560 minutes of observational data were obtained (mean ± standard deviation = 107.68 ± 21.22 minutes per focal individual). To investigate temporal consistency, which is a prerequisite of personality, focal data were split into two approximately equal non-overlapping phases (cf. Kluiver et al., 2022; Bhattacharjee et al., 2024a; Bhattacharjee et al., 2024b). The first phase consisted of data collected until April 2022, and the second phase included data collected between May and July 2022. All focal observations were filmed by a handheld Sony HDR-CX405 HD video camera.

**Table 1.** The rotated principal components or rhesus macaque personality traits with variable loadings and their percentages of variance, and percentages of attribute communalities of the behavioral variables.

| Variables | Sociability | Cautiousness | % Communalities |
| --- | --- | --- | --- |
| Look around | 0.87 |  | 72.99 |
| Approach | 0.81 |  | 71.99 |
| Avoid |  | 0.91 | 79.30 |
| Attention |  | 0.70 | 67.86 |
| **% Variance** | 38.19 | 33.58 |  |

Variable loadings of $>\pm 0.5$ are considered significant.

#### Novelty experiments

The experimental approach followed the standardized protocol for testing captive non-human primates (**see** Koski, 2011; Massen et al., 2013; Kluiver et al., 2022; Bhattacharjee et al., 2024a). We conducted tests involving novel objects, novel food items, food puzzles, and model predators to capture the ‘rare’ behaviors that are otherwise challenging to obtain through focal observations (e.g., exploration, persistence, boldness, etc.). For each experimental category, two different types of objects/food items/puzzles/predator models were used on different days to assess contextual consistencies in behavior (cf. Koski, 2011; Massen et al., 2013; Kluiver et al., 2022; Bhattacharjee et al., 2024a). Furthermore, for each type, except for the predator models, we provided multiple identical units (e.g., multiple novel objects of the same type) to avoid monopolization. All these experiments were conducted within the existing social group setting of the macaques in a pseudo-randomized order, ensuring that the same category tests were not repeated consecutively. We recorded all the experiments with a Sony HDR-CX405 HD video camera located at least 20 m from the experimental setups. Notably, focal observations were not conducted on the days of novelty experiments.

The novel objects were - (i) a hanging setup of glossy compact discs (CDs) and (ii) rubber toys. None of these objects contained food items inside. For the first object, a 230 cm long single string was threaded through 15-16 CDs (CD diameter = 12 cm). The two ends of the string were then tightly attached to existing structures (like wooden logs of park boundaries). We installed two such structures at distant locations approximately 120 cm above ground for the macaques to inspect and interact with (**Movie S1**). For the latter, locally made blue and red colored fish-shaped rubber toys were used (**Movie S2**). The rubber toys had a length of 7.6 cm and identical appearances. We provided five rubber toys to the macaques. The experimenter left the area after installing or placing the novel objects within a short duration (<10 seconds). The tests were filmed for 60 minutes after the placement of objects.

Novel food items were decided by observing macaques’ foraging activity (including provisioned items). We used gooseberries and kiwi as novel food items. Since kiwi fruits are larger than gooseberries, we made smaller pieces of kiwi fruits. Approximately 50 units or pieces of each food item were placed in an open area clearly visible to the macaques to avoid potential monopolization. After placing the food items, the experimenter left the area. We filmed the tests until all food items were eaten or for 60 minutes.

Two different types of food puzzles were used – (i) a wooden maze box and (ii) a rotating plastic bottle. We installed three wooden maze boxes (60 cm x 12 cm x 60 cm) approximately 80 cm above ground at distant locations in the study area. The boxes had plexiglass fronts with strategic openings where individuals could insert their fingers to move tomatoes (8-10 intact tomatoes) placed inside (**Movie S3**). The tomatoes could be retrieved by moving them in a specific direction through a larger opening. Like the wooden maze box, we installed three rotating bottles of 50 cm lengths at different locations. Each bottle was loaded with two handfuls of chickpeas and had 28 holes of 2.5 – 3 cm diameter on one side. The bottles were horizontally placed using tightly attached ∼ 200 cm strings, approximately 170 cm above ground. Originally, the holes were on the upper sides, but individuals could obtain the chickpeas through the holes by rotating and holding (often shaking) the bottles (**Movie S4**). The strings were attached in a way such that the bottles returned to their original positions (i.e., holes on the upper side) once individuals let go. After installing the puzzles, food items were loaded within 1 minute, and the experimenter left the area. We filmed the tests until all food items were retrieved or for 60 minutes.

For predator exposure experiments, we used snake and tiger models, both known to elicit antipredator responses in macaques (Maestripieri, 2010; Etting & Isbell, 2014; van Dijk et al., 2023). A rubber snake model (∼72 cm in length) with patterns resembling an Indian cobra was placed on the ground in a narrow path (**Movie S5**), which the study group frequently uses. The tiger model was a 108 cm-long plush toy in a sitting posture with patterns resembling a tiger (**Movie S6**). The tiger model was placed in the same position where the snake test took place. Unlike other experiments, each predator exposure experiment lasted for 30 minutes.

We intended to repeat the novelty experiments to test temporal consistencies in behavior; however, we found low participation rates and, therefore, decided not to repeat them.

### 2.3. Data coding

All behavioral and experimental videos were coded using ELAN (version 6.2, 2021). We followed the ethogram provided in **Table S1** to code behavioral data. Of the 87 behaviors described in the ethogram, only 35 behaviors were exhibited by at least 30% (16 individuals) of the study subjects. To avoid potential bias in analyses, we discarded the behaviors with low occurrences (i.e., exhibited by < 30% of the individuals). The 35 retained variables were corrected for the varying observation durations of the individuals, and their rates were calculated (seconds per minute for durational variables and frequency per minute for event behaviors). The standardized values of these variables were used in the analyses. From the novelty experiments, we coded the following parameters – latency to approach (novel objects, novel food items, food puzzles, and predator models), duration in proximity (novel objects, food puzzles, and predator models), number of approaches (predator models), duration of handling (novel objects), duration of manipulation (food puzzles), and eating (novel food items and food items from puzzles). See **Table S2** for definitions of these parameters. T.G. coded all data, and D.B. coded ∼ 6% of the total data (including observational and experimental data) to check for reliability. The reliability score was high (ICC (3,k) = 0.92, *p* < 0.001). As mentioned, we decided not to include experimental variables in the statistical analyses due to the low and inconsistent participation rates. Therefore, we proceeded with the observational data for the assessment of personality.

### 2.4. Statistical analyses

All statistical analyses were performed in R (version 4.3.0) (R Core Team, 2022). The repeatability of the 35 standardized behavioral variables between the two phases was assessed using a two-way mixed model intraclass correlation test (ICC (3,1)) (Lessells & Boag, 1987; McGraw & Wong, 1996). In the next step, we calculated the average values of the repeatable behavioral variables between the two phases and standardized them. A principal component analysis (PCA) was conducted using the average values of the repeatable variables. We checked whether the overall Kaiser-Meyer-Olkin sampling adequacy (MSA) value (cut-off value = 0.6) and attribute communality values (cut-off value = 60%) were high enough (Hadi et al., 2016). In case the values were below their cut-offs, the corresponding behavioral variables were identified, and a step-by-step process further reduced the number of variables until the cut-offs were crossed (cf. Kluiver et al., 2022; Bhattacharjee et al., 2024a). Once these criteria were met, a scree plot was generated based on an unrotated PCA, and the eigenvalue of each principal component (PC) was inspected. The number of PCs was decided depending on the eigenvalues (eigenvalue cut-off = ≥1) and by visual inspection of the scree plot. Further, a Bartlett’s sphericity test for correlation was conducted. As personality traits can correlate with each other and form behavioral syndromes, we used an oblique rotation technique (direct oblimin). The factor loadings ≥ 0.5 (positive and negative) were considered significant (Budaev, 2010), and from each PC, individual factor scores (synonymously personality scores) were obtained.

ICCs were run using the “ICC” function from the *psych* package (Revelle & Revelle, 2015). The PCA was performed using the “full_factor” function of *Radiant.multivariate* package (Nijs, 2021). We conducted linear models (LM) using the *glmmTMB* package to investigate the effect of sex on personality (Brooks et al., 2017). We constructed separate models (depending on the number of personality traits) with Gaussian error distributions with ‘identity’ link functions. Linear model diagnostics were investigated using the “testResiduals” function of the *DHARMa* package (Hartig, 2020). Null vs. full model comparisons were done using the “lrtest” function from the package *lmtest* (Zeileis & Hothorn, 2002). The significance value (α) was set at 0.05 for all statistical analyses.

### 2.5. Additional analyses on human-macaque interactions

Although the behavioral variables *contact provision* and *non-contact provision* were dropped due to low occurrences, we performed additional analyses as these variables are of great importance to synanthropic animals (Sengupta et al., 2015; Cox & Gaston, 2018; Balasubramaniam et al., 2020; Marty et al., 2020; Balasubramaniam et al., 2022). Based on data from both phases, we investigated which individuals obtained food through contact and non-contact provisioning. If an individual obtained food through both, we considered the response to be contact provisioning (securing food from human hands), as non-contact provisioning is considered the ‘default’ way of obtaining food (**Table 1**). We found that 15 individuals obtained food through contact provisioning, whereas the number was 17 for non-contact provisioning. We conducted Mann-Whitney U tests to compare the personality scores of individuals who obtained food through contact vs non-contact provisioning.

## 3. Results

### 3.1. Repeatability of behavioral variables

Of the 35 variables, we found nine to be repeatable (ICC value >0.2 and p < 0.05, **Table S3**). The ICC values ranged between 0.23 and 0.53, suggesting moderately low to relatively high repeatability. However, due to low overall MSA value (i.e., < 0.6) and inadequate attribute communality values (< 60%), the number of variables was reduced to four. The attribute communalities of the four remaining behavioral variables (*approach*, *attention*, *avoid*, and *look around*) ranged between ∼ 68% to 79%, with an overall MSA value of 0.61. All other assumptions of the PCA were fulfilled (Eigenvalues of selected PCs ≥ 1; Bartlett test: p < 0.001).

### 3.2. Personality traits

We found two personality traits in synanthropic rhesus macaques based on behavioral observations. These traits (PCs) cumulatively explained 71.77% of the variance. The labeling of traits was based on the variables loaded on them (**Figure 2**, **Table 1**). PC1 had two positively loaded variables: *look around* and *approach*. Look around is defined as an individual’s side-by-side head movements without any clear focus, whereas approach implies moving towards conspecifics in a neutral or affiliative manner, i.e., without displacing them. A lack of clear focus on conspecifics and a potential tendency to initiate affiliative social interactions prompted us to label PC1 as ‘sociability’. Sociability explained 38.19% of the variance of the data. Like sociability, PC2 included two positively loaded variables: *avoid* and *attention*. These behavioral variables included paying clear focus on conspecifics or within-group events. Avoid defines an individual’s tendency to redirect themselves when (certain) conspecifics are close by, while attention is focused on within-group conflicts and vocalizations, without moving head, often coupled with a frozen body posture. Accordingly, we labeled PC2 ‘cautiousness’. Cautiousness explained 33.58% of the data variance.

**Figure 2.**
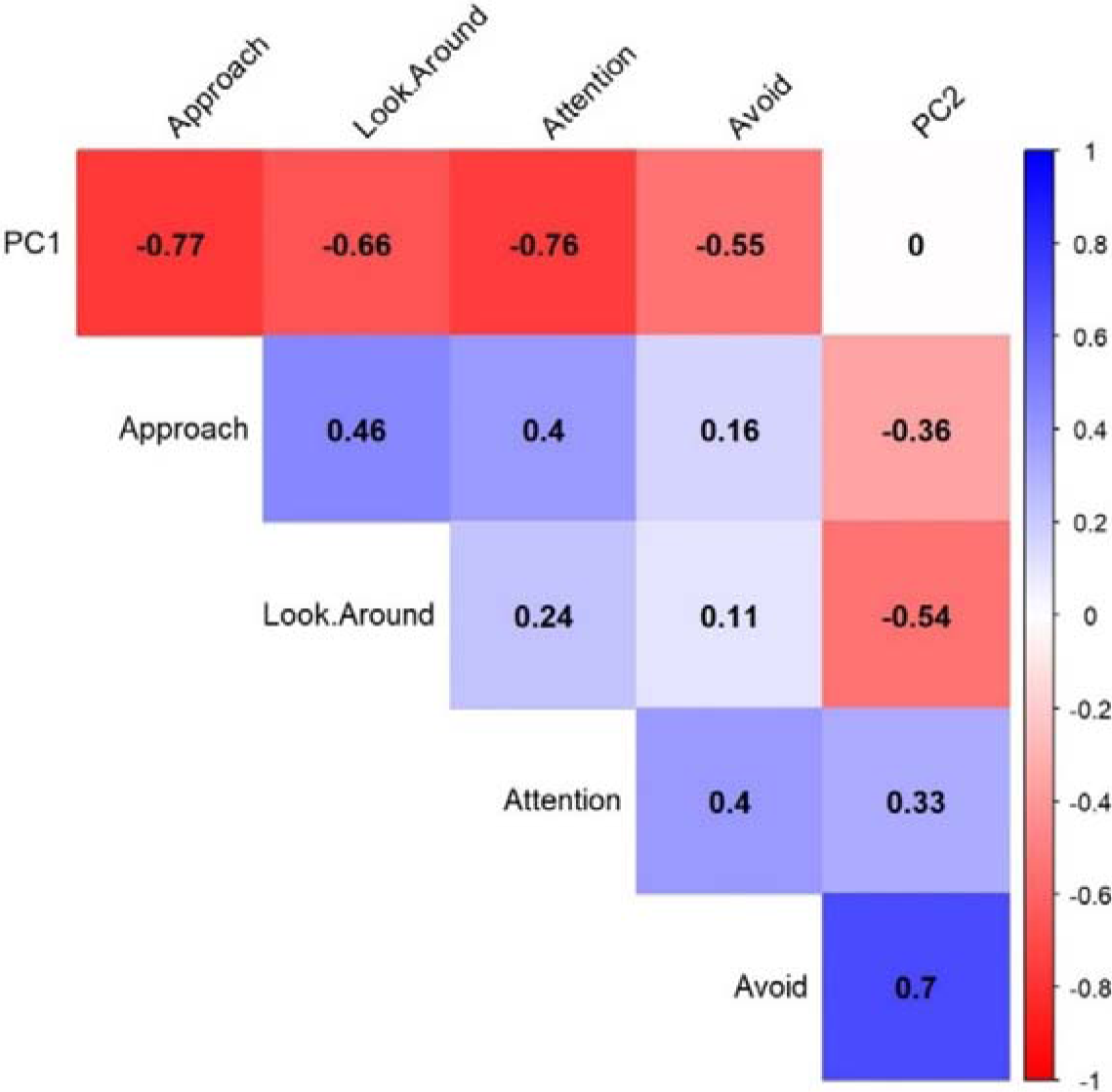
A correlation matrix plot of the behavioral variables and the principal components (PC1 and PC2).

### 3.3. Effect of sex on personality traits

We found an effect of sex on the sociability personality trait. Males were more sociable than females (LM: z = 2.43, Cohen’s d = 0.67, 95% CI [0.13, 1.22], p = 0.01). This full model differed from the null model that lacked the fixed effect of sex (p = 0.01). We did not find any effect of sex on cautiousness (LM: z = −0.934, Cohen’s d = 0.26, 95% CI [−0.29, 0.80], p = 0.35).

### 3.4. Response to novelty experiments

We found low participation in novelty experiments. Only six individuals (11.52% of the sample) participated and interacted with the novel object CDs. In contrast, only two individuals (3.84%) spent time near the rubber toys without handling them. Both the novel food item tests had only four individuals (7.69%). The food puzzles (wooden maze and rotating plastic bottle) had slightly higher participation rates than novel objects and novel food items. Nine individuals (17.30%) inspected and manipulated each of these puzzles. A total of ten individuals participated in the snake model (19.23%), whereas the tiger model had twelve participants (23.07%).

### 3.5. Human-macaque interactions during food provisioning

We found no difference in sociability trait scores between individuals who obtained food through contact and non-contact provisioning (Mann Whitney U test: U = 97, z = −0.89, p = 0.37, **Figure 3**). Unlike sociability, we found a significant difference in cautiousness trait scores between individuals who obtained food through contact and non-contact provisioning (Mann Whitney U test: U = 67, z = −2.07, p = 0.03, **Figure 3**). Individuals who obtained food through contact provisioning had higher cautiousness scores (i.e., more cautious) (mean ± SD: 0.74 ± 1.33) than individuals who obtained food through non-contact provisioning (−0.20 ± 0.50).

**Figure 3.**
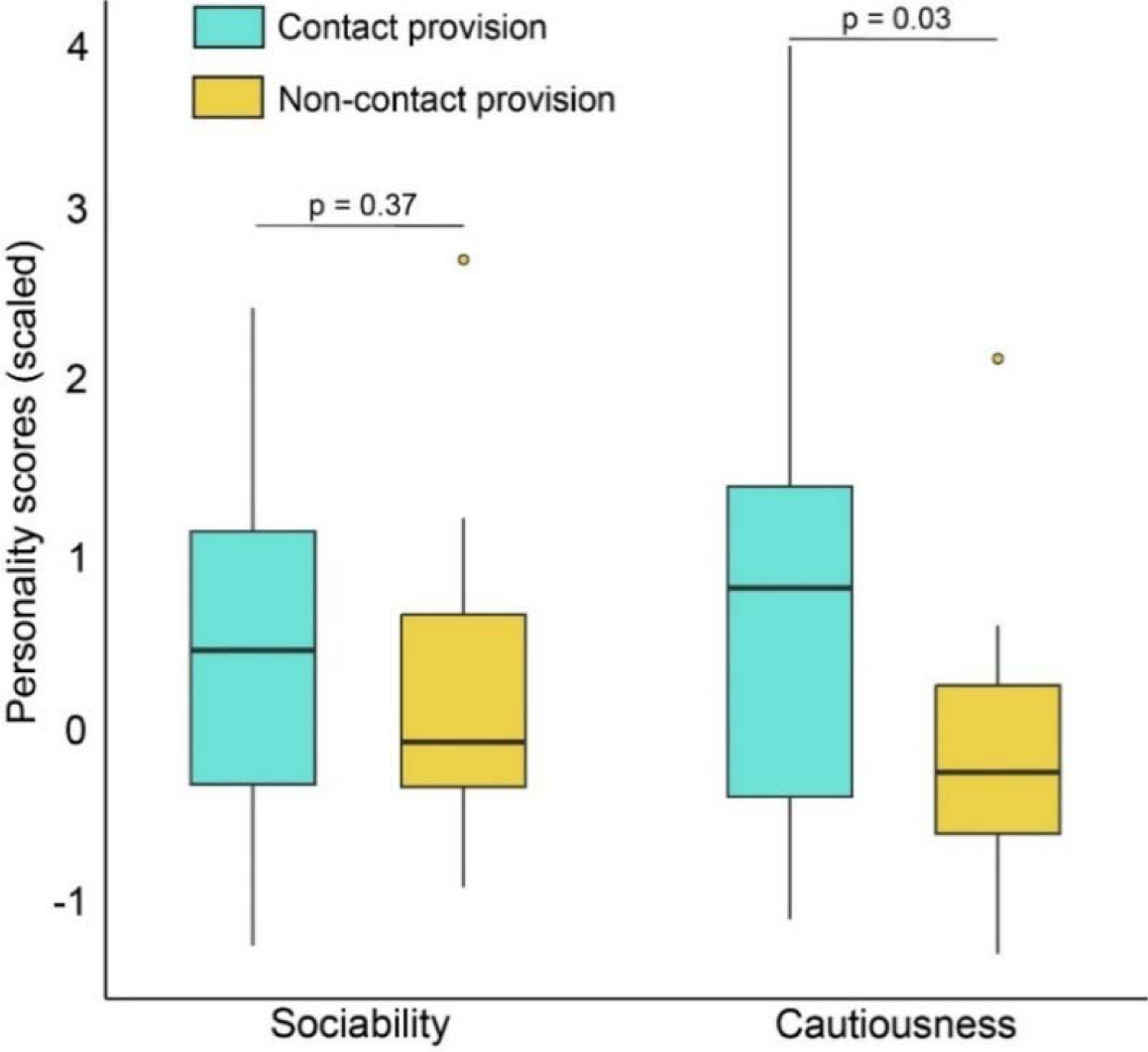
Personality traits and contact and non-contact food provision. Boxes indicate interquartile ranges, and whiskers represent the upper and lower limits of the data. The horizontal bars within the boxes represent the median values. Dots show outliers. Significance values (p-value) are provided.

## 4. Discussion

Animal personality research allows us to examine the ecological and evolutionary implications of biological traits and their variations. Yet, varying testing environments and different methodologies have made the task of assessing personalities notoriously challenging. Using a methodology standardized for testing captive NHPs, we intended to assess the personalities of synanthropic rhesus macaques in free-living conditions. Although we identified two traits based on behavioral observations and found partial associations with human-macaque interactions during provisioning, the extremely low participation of macaques hindered the successful implementation and completion of novelty experiments. Contrary to our prediction, males were more sociable than females. We discuss how observed personality traits and their variations can benefit a highly adapted NHP species living in human-dominated environments. We also recommend that future experimental studies should compare captive and non-captive populations to better understand the ecological validity of the experimental protocols.

### 4.1. Personality traits obtained through behavioral observations

The observed personality traits of sociability and cautiousness corroborate previous findings on rhesus macaque personalities that used similar observational approaches, albeit in partially different testing environments (Sussman et al., 2013; von Borell et al., 2016). However, the underlying behavioral variables of the personality constructs varied among these studies. This could be attributed to the different behavioral variables considered and the use of predefined categorizations in previous studies. For instance, the current study used a comprehensive ethogram of 110 behaviors (incl. passive behaviors), whereas others used ∼12 variables. Besides, we used a ‘bottom-up’ method instead of grouping multiple behaviors into predefined categories (such as physical aggression, submissiveness, etc.). While grouping behaviors into predefined categories may help tackle data of low representation, the approach is not free from inherent biases (such as lack of validity checks and presence of species/population-specific behaviors), which has been discussed previously (Uher, 2008; Freeman et al., 2013). Furthermore, irrespective of the methodological approach, labeling traits is still subjective, generating ambiguity on the interpretability of constructs among researchers. Nonetheless, we believe that animal personality research can significantly benefit from carrying out methodological comparisons.

Both personality traits reported in our study have elements of within-group-level social interactions and can be interpreted as ‘social personality traits’ (Koski, 2011). Like several other NHPs and generally gregarious animals, rhesus macaques show complex social dynamics (Sade, 2017), where social personality traits may play crucial roles. Consistency in sociality (i.e., an individual’s position in the social network) is linked to inter-individual variations in sociability in rhesus macaques (Brent et al., 2013), implying how sociability as a personality trait may shape group dynamics. Variations in sociability further hold the potential to explain group cohesion and cooperative interactions (Gartland et al., 2022, Bhattacharjee et al., 2024c). Rhesus macaques live in hierarchical matrilineal societies, where strong bonds among kin members, especially philopatric females, provide fitness benefits (Chapais, 1983). As more sociable individuals are more likely to engage in affiliative social interactions, higher sociable tendencies can help maintain these bonds (Gartland et al., 2022). Lower tendencies of and general variations in sociability could be explained by life-history trade-offs (pace-of-life syndrome, Réale et al., 2010), social niche specialization (Bergmüller & Taborsky, 2010), and state-behavior feedback (social niche hypothesis, Sih et al., 2015). Contrary to our prediction, we found that males were more sociable than females in our study group. This could be attributed to the necessity of males to form alliances (Higham & Maestripieri, 2010), potentially by interdependencies (Bhattacharjee et al., 2023). However, due to the low representation of males in our sample, conclusions should be drawn with caution. Variations in cautiousness, on the other hand, might be associated with the despotic social style of rhesus macaques, where within-group aggressive interactions are frequent (Thierry, 2007). Individuals can avoid potential conflicts by exercising increased caution. This could be achieved proximately by collecting spatiotemporal information on nearest neighbors (Sinha, 1998; Cameron & du Toit, 2005). Alternatively, by being less cautious, individuals may allocate their time and energy to other activities, like resting and feeding, or to events outside their social groups. In this study, the dominance rank relationships were not determined, which might have assisted in explaining how cautiousness could be linked to social style.

### 4.2. Challenges associated with experimental approach and proposed improvements

Replicating laboratory protocols of behavioral and cognitive experiments in the ‘wild’ is crucial to ensure ecological validity. While some studies successfully replicated laboratory protocols in the wild (Herborn et al., 2010), others encountered challenges (Carter et al., 2012; Schuppli et al., 2022). Our experimental approach was fairly inefficient, as evidenced by the low (average ∼ 13% for the eight tasks) and inconsistent participation rates of the individuals. Several factors, such as reduced neophobia in human-dominated environments, relatively large group size, conservative thresholds of coded variables, etc., might have contributed to this. We discuss these challenges in detail and propose improvements for future studies.

Living in highly enriched human-dominated environments makes animals less neophobic toward human artifacts (Griffin et al., 2017; Bhattacharjee et al., 2024d). We suspect the ‘novel’ objects were not novel enough to elicit behavioral responses, resulting in the individuals’ lack of engagement. Similar patterns were also observed in semi free-ranging Japanese macaques (*Macaca fuscata*), where individuals lacked a tendency to interact with novel objects (Personal observations, J.J.M.M., D.B., 2021-2023). Here, as opposed to adults, we found juveniles inspected and interacted with novel objects more, which could be explained by their relatively higher tendency to explore and innovate (Kendal et al., 2005). However, we only focused on adult personalities in this study. The primary reason for low participation in novel food item tasks could be associated with the fact that our study group is regularly provisioned with ‘highly rewarding’ food items, like apples, bananas, and peanuts. Provisioning may affect animals’ exploration tendencies (Berman & Li, 2002; Sengupta et al., 2015). Nonetheless, relatively larger food items (such as dragon fruits and durians, if they are potentially novel) could help generate more attention from synanthropic animals. We also think conducting the novel food item tests in multiple areas simultaneously would be beneficial in capturing the behavioral responses of a relatively large population, like in our study. In contrast, the food puzzle tests garnered relatively higher participation. However, the fast-depleting nature of food items in the puzzles potentially led to less overall involvement. A relatively costly alternative would be remotely controlled food dispensers, as such devices have proved fruitful in capturing animals’ behavioral and cognitive responses in the wild (Rosati, 2022). Additionally, ‘impossible food tasks’ could be used to examine traits like persistence. Finally, for the predator exposure experiments, we coded variables (especially the distances) like the captive conditions (cf. Kluiver et al., 2022; van Dijk et al., 2023; Bhattacharjee et al., 2024a). While we found ∼20% participation in the predator experiments, more data could have been obtained by making the variables less conservative, i.e., by increasing the radius for approach, proximity, etc. Besides, life-sized predator models could elicit more intense antipredator behaviors (cf. Kluiver et al., 2022)

### 4.3. Implications of macaque personality on human-macaque interactions

Our additional analyses revealed an association between personality and human-macaque interactions during provisioning. While sociability did not differ, individuals obtaining food through contact provisioning were more cautious than individuals who obtained food through non-contact provisioning. Notably, cautiousness as a personality trait was measured at the level of within-group interactions. Therefore, cautiousness for conspecifics does not necessarily translate into identical behavioral responses while interacting with humans. Instead, this response could indicate higher risk-taking or bolder traits in more cautious individuals, thus potentially suggesting a behavioral syndrome. Besides, if more cautious individuals are the low-ranking individuals in a group, obtaining food by making contact with humans can benefit them. It can help secure provisioned food items using an alternate yet effective strategy by not waiting for the food items to be placed on the ground, where more dominant individuals can exhibit aggression. In a comparative study using despotic Japanese macaques, slightly less despotic barbary macaques (*Macaca sylvanus*), and more egalitarian moor macaques (*Macaca maura*), researchers found that subordinates in the more despotic societies use alternate ‘opportunistic’ strategies to maximize food intake (Gomez-Melara et al., 2021). More systematic observations in the future would be valuable for a nuanced understanding of how variations in traits are linked to intra-and inter-specific interactions.

From a ‘One health’ perspective, provisioning can have severe consequences on human-animal coexistence. The act of provisioning affects the behavioral dynamics of animals in human-dominated environments (and beyond) and can also foster the transmission of zoonotic diseases (Strandin et al., 2018; Shutt & Lees, 2021). Growing human-macaque conflicts in Asia and Africa, as well as macaques’ tendencies to ‘bargain’ in several Asian countries, are classic examples of ‘provisioning gone wrong’ (Riley, 2007; Fuentes et al., 2008; Radhakrishna et al., 2012; Priston & McLennan, 2012). Unfortunately, how these behavioral dynamics relate to personalities is poorly understood. Therefore, more studies on synanthropic NHP personalities would be valuable for managing populations and mitigating conflicts between humans and NHPs.

## Abbreviations

NHP: Non-human primates
KNR: Kamla Nehru Ridge Biodiversity Park
PCA: Principal Component Analysis
UGC: University Grants Commission
MSCA-IF: Marie Sklodowska-Curie Actions Individual Fellowship

## Author contributions

Tania Gill: Investigation, data curation, formal analysis, writing— original draft; Anshul Gautam: investigation, data curation; Jorg J.M. Massen: Conceptualization, methodology, resources, writing—review and editing. Debottam Bhattacharjee: Conceptualization, formal analysis, supervision, resources, visualization, writing—original draft, writing—review and editing. All authors gave their final approval for publication and agreed to be held accountable for the work performed therein.

## Institutional Review Board Statement

Ethical approval for the study was obtained from the Department of Forests and Wildlife, Government of NCT of Delhi, India (Approval No.: F.No.CF/LC/105/07/HQ/PartFile-/833-37). All experiments were non-invasive, and the participation of animals was completely voluntary. We adhered to the ethical guidelines of the American Society of Primatologists for the study.

## Data Availability Statement

All data generated in the study and the code for analyses (R-script) will be made open access upon publication.

## Acknowledgments

We are grateful to Nisheeth Saxena, the Chief Conservator of Forests/Chief Wildlife Warden, Department of Forests and Wildlife, Government of NCT of Delhi, India, for granting permission to study free-ranging rhesus macaques at the Kamla Nehru Ridge Biodiversity Park, New Delhi, India. We want to thank Pankhuri Singhal, Tarangini Chutia, and Akanksh Pandey for helping with the data collection and organization. We acknowledge Eva van Dijk for her assistance in the data organization. We also thank Avitoli G. Zhimo and H.N. Kumara for guiding T.G. in obtaining institutional support during the study. T.G. was supported by a senior research fellowship funded by the University Grants Commission, Government of India (reference number 6894, June 2015). D.B. was funded by the European Union’s Horizon 2020 research and innovation program, Marie Sklodowska-Curie Actions (grant number H2020-MSCA-IF-2019-893016).

