## Supplementary Information for "Personality assessment of synanthropic rhesus macaques: implications and challenges"

by

Taniya Gill, Anshul Gautam, Jorg J.M. Massen**,** and Debottam Bhattacharjee

| **Table of contents** | **Page no.** |
| --- | --- |
| Figure S1. Rhesus macaques feeding on provisioned food items | 02 |
| Table S1. Ethogram used in the study | 03 - 09 |
| Table S2. Variables coded from novelty experiments | 10 |
| Table S3. Intra-class correlation test results | 11 |
| Supplementary media files | 12 |

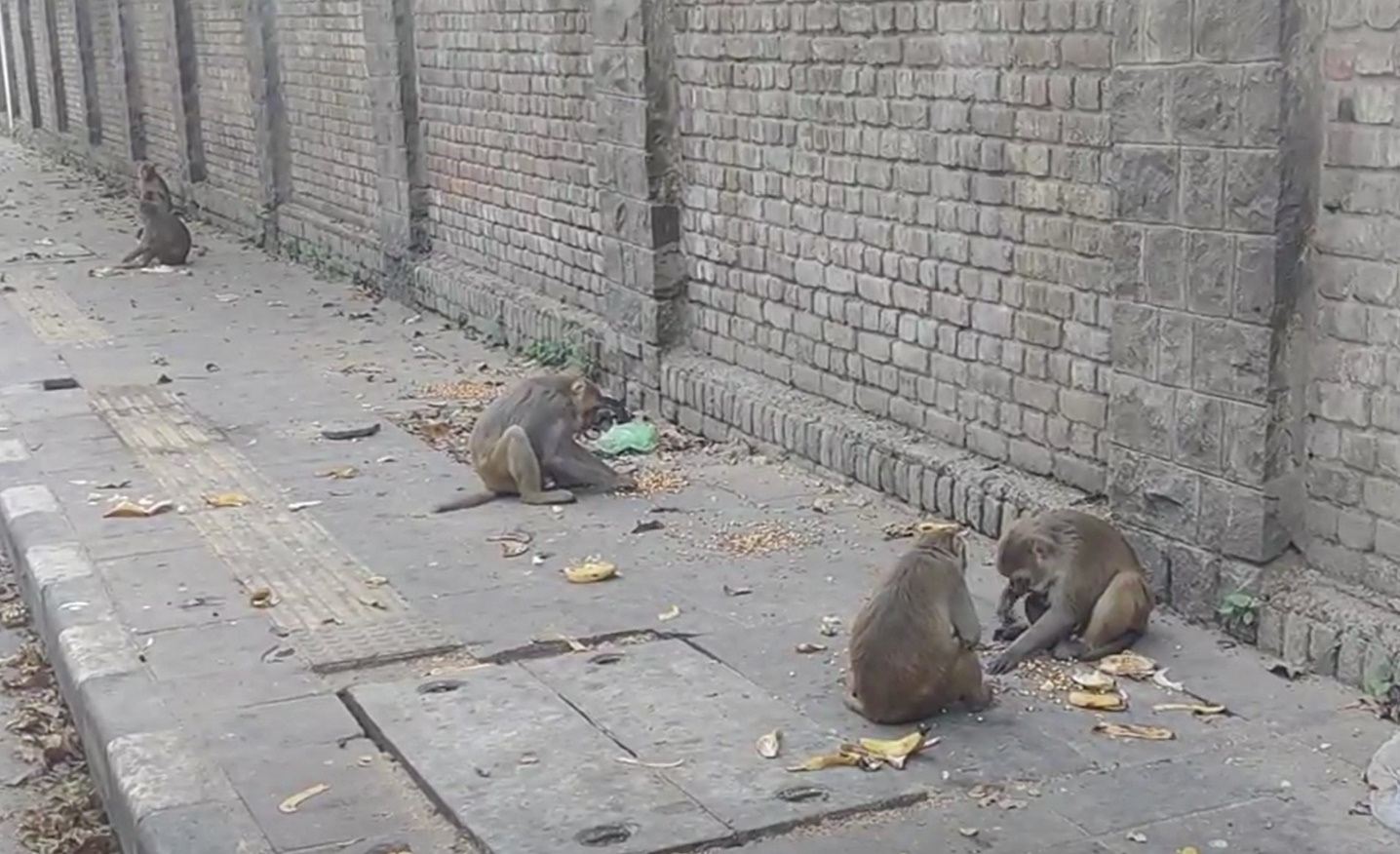

**Figure S1. The rhesus macaques feeding on provisioned food items at the Kamla Nehru Ridge Biodiversity Park.** [Photo credit: Taniya Gill, 2022]

**Table S1. Ethogram used for behavioral observations and coding. Names of behaviors, types, and their descriptions are summarized.**

| **Behavior** | **Type** | **Description** |
| --- | --- | --- |
| *Activity* | | |
| Forage | Durational behavior | Focal moves slowly/ sitting/ standing while looking for food on the ground |
| Active forage | Durational behavior | Focal actively eats/ handles food items |
| Regurgitation | Durational behavior | Chewing of previously ingested food (often from cheek pouches) without actively handling food items |
| Sit | Durational behavior | Focal rests in a posture supported by the buttocks or thighs |
| Travel | Durational behavior | Focal walks or runs around |
| Stand | Durational behavior | Focal stands on four legs |
| Bipedal stand | Durational behavior | Focal stands on hindlegs |
| Curl up | Durational behavior | Focal sits hunched forward with arms and legs tight at the body core and head tilted towards the chest. |
| Climb up/down | Durational behavior | Focal moves in any direction on a tree, building, or construction |
| Jump | Event behavior | Focal jumps in the air, but not towards another individual |
| Lie down | Durational behavior | Focal is lying down, either on the side, belly, or back |
| Bipedal walk | Durational behavior | Focal walks standing up on both hindlegs |
| Object play | Durational behavior | Focal uses objects in movements that serve no obvious, immediate purpose |
| Object manipulation | Durational behavior | Focal handles or manipulates non-food objects with a clear purpose |
| Reach | Event behavior | Focal reaches towards an object out of reach or without a clear purpose involving grabbing-like behavior |
| Stretch | Event behavior | Focal stretches its body by keeping the hindlegs out, walking the front legs forward, and slightly dropping the pelvis, which might include bulging the back. |
| Scream | Event behavior | Focal utter high-pitched, intense vocalization. Can be continuous when the focal screams every 10 seconds |
| Pass by | Event behavior | Focal enters a radius of 1 meter around another individual and leaves this radius within the same movement |
| Defecate | Event behavior | Focal releases feces, usually in a sitting or squatting position |
| Urinate | Event behavior | Focal releases urine, usually in a sitting or squatting position |
| Steal food/object | Event behavior | Grabbing a piece of food or an object (e.g. toy) from another individual, possibly accompanied by resistance from the other individual |
| Startle | Event behavior | Focal shows a frightened response (frozen body posture with attention towards stimuli) due to a sudden change in the environment (e.g., loud sound, other members making moves, or loud noises with human artifacts) |
| Drink water | Durational behavior | Focal visibly drinks water by bending its body and making contact with any water resource |
| *Sexual behavior* | | |
| Sexual Present | Event behavior | Focal displays its rump and/or anogenital region to another individual by bowing forward and raising the hindquarters |
| Sexual Mount | Event behavior | Focal climbs on another adult individual with its hands on the receiver's back or rump. Sexual mounting might involve thrusts. Only noted when the focal is in consortship with the other individual |
| Masturbate | Durational behavior | Focal rubs own genitalia with fingers, on objects, or on the ground |
| Hold bottom | Event behavior | Focal holds another buttocks or hips from behind for a short while |
| Inspection | Event behavior | Focal pulling the tail of a female aside and visually inspecting the vulva and anus can also include sniffing |
| Ground hit | Event behavior | Focal hits or wipes the floor with the hand |
| Groom rejection | Event behavior | Focal rejects being groomed by another individual by grabbing or pushing away the arm of the other individual |
| *Social behavior* | | |
| Approach | Event behavior | Focal moves towards a conspecific with clear focus and ends up within a 1-meter radius of the conspecific |
| Touch | Event behavior | Focal makes gentle and short body contact with another individual |
| Contact sit | Durational behavior | Focal is sitting next to another individual, and body parts touch (excluding limbs) |
| Contact lie down | Durational behavior | Focal is lying next to another individual, and body parts touch (excluding limbs) |
| Proximity | Durational behavior | Focal is within 1 meter of another individual |
| Change position | Event behavior | Adjusting position without leaving the immediate surroundings/the contact sit. Moving less than 1 meter |
| Co-feed | Durational behavior | Focal is actively foraging with another actively foraging individual (both from the same source) while being within 1 meter from each other |
| Groom | Durational behavior | Focal touches and strokes another individual's fur with one or both hands, accompanied by periodic hand contact with its own mouth. Focal pays close attention to the recipient’s fur |
| Allogroom | Durational behavior | Focal is involved in a multiple grooming interaction where the focal is being groomed and grooms at the same time |
| Autogroom | Durational behavior | Focal grooms oneself or closely inspects skin, nails, hands, toes, or other body part |
| Groom present | Event behavior | Focal approaches (or is approached by) another individual and immediately lies down when in proximity to the other individual, |
| Social play | Event behavior | Focal displays behaviors of physical aggression (e.g. hit, grab, or bite) with other individuals without vocalizations and with less intensity than regular physical aggression |
| Vocalization | Event behavior | Focal utters single vocalization of regular intensity and pitch (i.e. no scream, grunt, or cry for support) |
| Travel together | Durational behavior | Focal travels parallel to another individual. Distance between the individuals is less than 3 meters |
| Carry | Durational behavior | Focal carries a juvenile/ infant on the back/belly |
| Nurse | Durational behavior | Focal holds infant close to breast in order to breastfeed |
| Auto nursing | Durational behavior | Focal breastfeeds herself |
| Follow | Event behavior | Focal walks behind another individual in the same direction and stays within a radius of 5 meters |
| *Dominance and submission* | | |
| Leave | Event behavior | Focal walks away from another individual within a radius of 1 meter or after being approached by/lunged at by another individual |
| Mock leave | Event behavior | Focal walks away from another individual while turning around at least once to lunge/scream/threaten the other individual |
| Displace | Event behavior | Focal approaches another individual who then leaves, after which the focal sits within a 1m radius where the other animal used to stay |
| Cover | Event behavior | A lateral flexion of the spine, away from another individual |
| Avoid | Event behavior | Focal changes its direction while walking or sitting spot to a distance of at least 3 meters from another individual that approaches the focal. The attention of the focal is on the approaching individual |
| Avoidance of eye-contact | Event behavior | Focal looks around undirected when another individual focuses their attention on the focal. Can be combined with autogrooming or playing on the ground |
| Freeze | Event behavior | Focal stops displaying body movements and looks to the ground while another individual passes by. Duration is a maximum of 20 seconds. |
| Social present | Event behavior | Focal displays its rump and/or anogenital region to another individual by bowing forward and raising the hindquarters. Focal might also sit in front of a potential partner and displays the back towards another individual. Could involve hitting the ground. Only noted outside consortship |
| Social Mount | Event behavior | Focal climbs on another adult individual with its hands on the receiver's back or rump. Only noted outside consortship |
| *Conflict and aggression* | | |
| Cry for support | Event behavior | During or after an aggressive encounter, the focal makes a high-pitched vocalization accompanied by hectic looking and searching for another individual. Focal has a tense body posture |
| Aggressive intervention | Event behavior | Focal approaches the center of a conflict and displays aggressive behaviors (e.g., lunge, threat, physical attacks, and chase). Focal was not involved at the start of the conflict |
| Leave Conflict | Event behavior | Focal leaves conflict, which is happening within 3 meters |
| Support | Event behavior | Focal actively approaches another individual who is in a conflict, and together, they chase away one opponent. When a juvenile is supported, the juvenile might leave the conflict while the focal is chasing |
| Stare | Event behavior | Focal looks intensively at another individual with a straight back and raised eyebrows |
| Chase | Event behavior | Focal runs after another individual for at least 3 meters |
| Lunge | Event behavior | Focal jumps a maximum of two body lengths forward toward another individual |
| Grab | Event behavior | Focal holds body parts of another individual with one or two hands for a few seconds |
| Hit | Event behavior | Focal slaps another individual by hand |
| Bite | Event behavior | Focal places its teeth on the body of another individual and closes its jaws |
| Redirection | Event behavior | Focal that was involved in a conflict displays the aggression to another, not previously involved individual within 10 seconds after the conflict (e.g. threats and/or physical aggression/chase/lunge) |
| Tree shake | Event behavior | Focal holds a tree or object and shakes its body intensively |
| Flee | Event behavior | Focal runs away from another individual after being approached/chased/lunged by another individual |
| Mock flee | Event behavior | Focal runs away from another individual while turning around at least once to lunge/scream/threaten the other individual |
| Push | Event behavior | pushing another individual away |
| *Tension* | | |
| Scratch | Event behavior | Focal uses finger, hand, or foot to rake across own skin |
| Yawn | Event behavior | Focal opens mouth wide and inhales intensely, which can be seen by the expansion of the chest |
| Body shake | Event behavior | Focal rapidly turns the whole body in at least two different directions |
| Vigilance | Event behavior | Focal has a tense body posture and looks around with hasty movements of head and/or eyes without an imminent reason. Might be accompanied by standing on the hind legs |
| Avoid conflict | Event behavior | Focal focuses its attention on a conflict without approaching the conflict and moves away in the opposite direction |
| Attention | Event behavior | Focal looks towards a specific situation (e.g., conflict, disturbance, or vocalization), individual, or object with a clear focus without moving its head and eyes and with a frozen body posture |
| Look around | Event behavior | Focal moves its head in at least three different directions without a clear Focus. Movements are not up and down |
| *Facial expressions* | | |
| Raise brow | Event behavior | Focal raises eyebrows at another individual; the head is slightly lifted and angled. Usually displayed when an individual is approaching another |
| Fear grimace | Event behavior | Focal pulls the corners of the lips back while slightly opening the mouth; teeth are visible. Often shown when receiving aggression |
| Open mouth threat | Event behavior | Focal opens his mouth for a while, directed at the receiver if aggression. Chin often points forwards |
| Lip smacking | Event behavior | Focal opens and closes mouth rapidly, pressing the lips closely together |
| *Vocalizations* | | |
| Soft grunt | Event behavior | A grunting sound, often emitted in series and uttered in affiliative interactions and contacts |
| Low coo | Event behavior | A brief and rather nasal sound of low frequency. This is a contact call exchanged by individuals in proximity |
| *Human-macaque interactions during provisioning* | | |
| Contact provision | Event behavior | Focal makes physical contact and obtains food item(s) from human hand during provisioning |
| Non-contact provision | Event behavior | Focal obtains food item(s) from the ground during provisioning |

**Table S2. Description of variables coded from novelty experiments to assess personality of rhesus macaques.**

| **Experiment** | **Variable** | **Description** |
| --- | --- | --- |
| **Predator model** | Latency to approach | Time taken by an individual to enter the 2-meter radius of the predator model for the first time from an initial distance of 5 meters |
|  | Time in proximity | Cumulative time an individual spends in proximity (< 2 meters) to the predator model |
|  | Number of approaches | The total number of times an individual approaches the predator model |
| **Novel object** | Latency to approach | Time taken by an individual to enter the 1-meter radius of the novel object for the first time from an initial distance of 5 meters |
|  | Time in proximity | Cumulative time an individual spends in proximity (< 1 meter) to the novel object |
|  | Time handling | Cumulative time an individual spends touching/holding (< 1 meter) the novel object |
| **Novel food** | Latency to approach | Time taken by an individual to enter a 1-meter radius of the novel food for the first time from an initial distance of five meters |
|  | Eat | Whether an individual eats the novel food. Noted as yes or no. At least one instance of visible eating |
| **Food puzzle** | Latency to approach | Time taken by an individual to enter the 1-meter radius of the food puzzle for the first time from an initial distance of five meters |
|  | Time in proximity | Cumulative time an individual spends in proximity (< 1 meter) to the food puzzle |
|  | Time manipulating | Cumulative time an individual spends manipulating (touching/ handling/licking/biting) food puzzles |
|  | Success in obtaining rewards | Whether an individual successfully obtains a reward from a puzzle. Noted as yes or no |

**Table S3. Intra-class correlation test results showing the repeatability values (coefficients, F- and p-values) of the behavioral variables for personality assessment.**

| **Behavioral variable** | **ICC** | **F-value** | **p-value** |
| --- | --- | --- | --- |
| **Approach** | **0.229822738** | 1.596804785 | **0.048913523** |
| Approach passive | -0.094303395 | 0.827646711 | 0.749102341 |
| **Attention** | **0.531591627** | 3.269778503 | **< 0.001** |
| **Autogroom** | **0.291416581** | 1.822532883 | **0.017123713** |
| **Avoid** | **0.258665303** | 1.697836765 | **0.030729722** |
| Avoid passive | -0.157760297 | 0.727473299 | 0.870383983 |
| Change position | -0.009285638 | 0.981599583 | 0.526307067 |
| Climb | -0.119768293 | 0.786083793 | 0.803507282 |
| Co-feed | 0.064851516 | 1.138697794 | 0.322272187 |
| **Contact lie down** | **0.333200703** | 1.99940328 | **0.007384198** |
| Contact sit | -0.020543611 | 0.959739866 | 0.558042779 |
| Curl up | -0.001113726 | 0.997775026 | 0.503157478 |
| Drink water | 0.097321852 | 1.215629131 | 0.244070847 |
| Flee | 0.169585477 | 1.408435721 | 0.11237808 |
| Flee passive | -0.043364778 | 0.916875136 | 0.621079611 |
| Forage | 0.104190753 | 1.232618167 | 0.228905162 |
| Groom | 0.147950427 | 1.347281266 | 0.145200201 |
| Groom passive | -0.052747436 | 0.899790901 | 0.646211304 |
| Groom present | -0.118191927 | 0.788601716 | 0.800355758 |
| Jump | 0.032153493 | 1.066443372 | 0.409606125 |
| Lie down | 0.022857692 | 1.046784776 | 0.435471946 |
| **Look around** | **0.243787449** | 1.644759066 | **0.039277979** |
| Lunge | 0.156613796 | 1.371392833 | 0.13137695 |
| **Open mouth threat** | **0.231818183** | 1.603550302 | **0.047434204** |
| Pass by | 0.056942256 | 1.120760911 | 0.342742499 |
| Pass by passive | 0.140640834 | 1.327315608 | 0.157576144 |
| **Proximity** | **0.248406414** | 1.661012597 | **0.036443967** |
| **Scratch** | **0.256360726** | 1.689476028 | **0.031946194** |
| Sit | -0.158004739 | 0.727108649 | 0.870757364 |
| Stand | 0.060106985 | 1.127901759 | 0.334493738 |
| Startle | -0.041275171 | 0.920721876 | 0.615409991 |
| Travel | 0.151555951 | 1.357256206 | 0.139336096 |
| Travel together | -0.069416797 | 0.870178219 | 0.689314399 |
| Vigilance | 0.113103122 | 1.255053604 | 0.210017318 |
| Yawn | 0.184028988 | 1.451067464 | 0.093567328 |

ICC values > 0.2 and p < 0.05 suggest repeatable behavioral variables and are shown in bold-type fonts

**Supplementary media files**

**Movie S1.** Rhesus macaques interacting with a hanging setup of glossy compact discs.

**Movie S2.** Rhesus macaques handling a rubber toy.

**Movie S3.** Rhesus macaques manipulating a wooden puzzle box with tomatoes.

**Movie S4.** Rhesus macaques manipulating a rotating bottle with chickpeas.

**Movie S5.** Rhesus macaques inspecting a snake model.

**Movie S6.** Rhesus macaques inspecting a tiger model.
